# Acoustic properties across the human skull

**DOI:** 10.1101/2021.04.22.440927

**Authors:** Thomas Riis, Taylor Webb, Jan Kubanek

## Abstract

Transcranial ultrasound is emerging as a noninvasive tool for targeted treatments of brain disorders. Transcranial ultrasound has been used for remotely mediated surgeries, transient opening of the blood-brain barrier, local drug delivery, and neuromodulation. However, all applications have been limited by the severe attenuation and phase distortion of ultrasound by the skull. Here, we characterized the dependence of the aberrations on specific anatomical segments of the skull. In particular, we measured ultrasound propagation properties throughout the perimeter of intact human skulls at 500 kHz. We found that the parietal bone provides substantially higher transmission (average pressure transmission 31±7%) and smaller phase distortion (242±44 degrees) than frontal (13±2%, 425±47 degrees) and occipital bone regions (16±4%, 416±35 degrees). In addition, we found that across skull regions, transmission strongly anti-correlated (*R* = −0.79) and phase distortion correlated (*R* = 0.85) with skull thickness. This information guides the design, positioning, and skull correction functionality of next-generation devices for effective, safe, and reproducible transcranial focused ultrasound therapies.

## 1. Introduction

Transcranial focused ultrasound offers incisionless and targeted treatment options for disorders of brain function [1–3]. At high intensities, ultrasound has been used to lesion malfunctioning or diseased deep brain targets [4, 5]. At low intensities, ultrasound can be used to deliver large drugs, genes, or stem cells across the blood-brain barrier [6–8]; release drugs in specific brain regions without affecting the blood-brain barrier [9–11]; and to modulate neural activity in a transient [12–15] or sustained [16–22] fashion.

The clinical utility of these applications has been impeded by the severe aberrations of ultrasound by the human skull. In the surgical applications, the highly variable attenuation and phase distortions can leave a substantial proportion of patients untreated [23]. The skull aberrations have limited the predictability of the ultrasound magnitude delivered into target [24] and so have also impeded the translation of low-intensity applications. A tight control of the delivered dose is particularly important for ultrasound-mediated opening of the blood-brain barrier, in which small changes in the ultrasound pressure at target can lead to vast differences in the scale of the blood-brain barrier disruption [6, 25]. The skull aberrations have also curbed effective and reproducible applications of ultrasonic neuromodulation [26, 27] and local drug release, which require a well-defined ultrasound dose [10, 28].

To maximize ultrasound penetration through the skull and to maximize the predictability of the delivered dose, ultrasound should be applied through skull segments that cause the least amount of attenuation and phase distortion. In diagnostic applications, small imaging probes can be applied through the temporal window, which has a relatively low ultrasound attenuation [29–32]. However, therapeutic applications often require large apertures or arrays with many elements for focal delivery of considerable amount of energy [4, 16, 17, 26, 33–36]. Any future design that maximizes ultrasound penetration and the predictability of the delivered dose should take into account the dependence of the aberrations on specific anatomical regions of the skull.

This information is currently incomplete. Existing studies have provided insights into inter-subject variability of acoustic properties [29, 37–40], estimates of average attenuation and phase distortions [29, 37–46], as well as approaches on how these aberrations may be compensated for [4, 47–53]. However, acoustic measurements have only been provided for discrete sets of chosen samples or skull flaps [29, 37–41]. Consequently, there is no systematic assessment of acoustic propagation properties within single intact skulls as a function of anatomical location.

To address this, we devised a setup that allowed us to measure ultrasound propagation properties throughout the perimeter of intact human skulls. We complemented these acoustic measurements with caliper measurement of the corresponding skull thickness. The resulting data quantify the transmission and phase distortion through anatomically defined segments of the skull, and show how these variables depend on the skull thickness. This information guides the design and placement of future devices for effective applications of transcranial ultrasound in the clinics.

## 2. Materials and Methods

### 2.1. Subjects

Three *ex-vivo* skulls were used in this study (Skull 1: male, 84 years; Skull 2: male, 61 years; Skull 3: female 68 years). The skulls were obtained under a research agreement from Skulls Unlimited (Oklahoma City, OK). An opening was made at the bottom of each skull to enable skull rotation around a receiving transducer positioned inside the skull.

Each skull was degassed overnight in a deionized water. Following the degassing, the skull was transferred, within the degassed water, into an experimental tank filled with continuously degassed water (AIMS III system with AQUAS-10 Water Conditioner, Onda). The water conditioner (AQUAS-10) treats water for ultrasound measurements in compliance with IEC 62781. The conditioner degasses water to remove undesired bubbles, removes suspended particles and biological contaminants, and deionizes water. The dissolved oxygen is between 2.0-2.5 PPM during continuous operation, according to measurements provided by the manufacturer (Onda). In comparison, tap water contains about 10.5 PPM of dissolved oxygen.

The skull was held in place by a pair of thin neodymium rare earth magnets (550lbs lift, 2.5-inch diameter, Neosmuk), one positioned below the skull and one above, at the center of the sagittal suture. The magnets allowed us to firmly hold and rotate each skull without having to perturb its surface.

### 2.2. Coordinates

Prior to degassing, the through-transmit plane was established using 4 markers that were chosen such as to avoid frontal and sphenoidal sinuses and to maintain perpendicular ultrasound propagation. Specifically, a frontal marker was positioned 49 mm above the center of the nasion. Two parietal markers, one on the left and one on the right, were made 17 mm above the squamous suture and at the widest point of the skull. The final, occipital marker was made approximately 17 mm above the center of the inion. An angular positioner assembly (AP02-S, Onda) was aligned with the 4 markers in the following way: the frontal marker corresponded to 0 degrees, the right parietal marker to 90 degrees, the occipital marker to 180 degrees, and the left parietal marker to 270 degrees (Fig. 1b).

**Figure 1:**
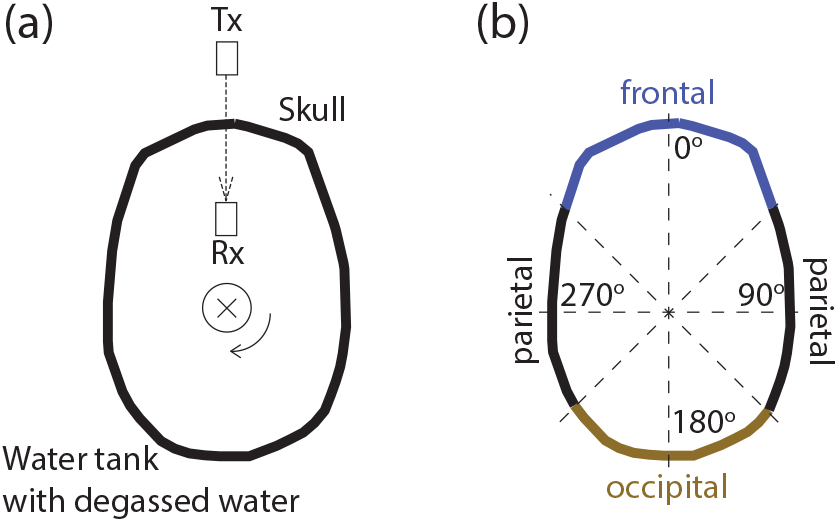
Through-transmit measurements across intact skulls. (a) Top view of the setup. Degassed and hydrated *ex-vivo* skulls were held by a robotic arm and were secured to the skull using two opposing magnets positioned at the center of the sagittal suture (the thin circle shows the position of the magnets). This robotic arm, connected to the magnet at the top of the skull, allowed us to electronically rotate the skull and so collect through-transmit measurements over individual segments of the skull within the imaging plane. The through-transmit measurements were achieved using a transmitting (Tx) and a receiving (Rx) transducer facing each other at a distance of 100 mm. The direction of ultrasound transmission is indicated by the dashed arrow. The through-transmit measurements were acquired at each rotation step of 1 degree. (b) Parameterization of the skull bone into parietal (45–135 and 225–315 degrees), occipital (135–225 degrees), and frontal (315–45 degrees) regions.

**Figure 2:**
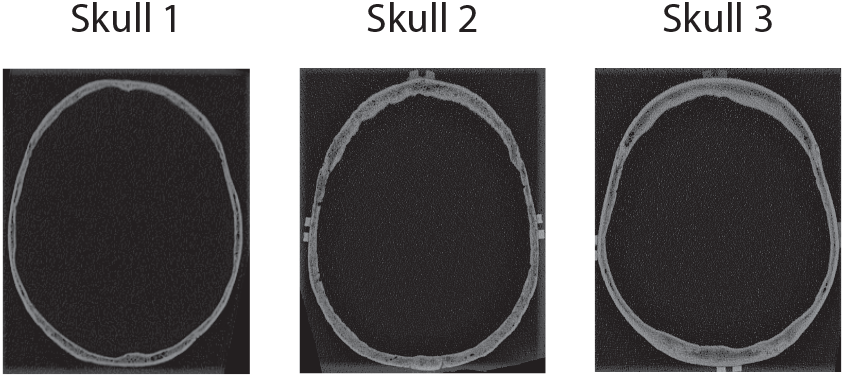
Subjects. CT scans for the three *ex-vivo* skulls used in this study. The images were taken at the through-transmit plane.

### 2.3. Skull thickness measurements

To measure skull thickness, the 4 markers were connected with a line. A precision caliper gage (Fowler 54-554-630, 0.1 mm accuracy) was used to measure the thickness of the skull in 3 mm steps. Each measurement was repeated 3 times; the resulting values were averaged together. Only rarely was there a difference of more than 0.1 mm between the 3 measurements.

### 2.4. Through-transmit setup

Two transducers operating at 500 kHz center frequency (V301-SU, Olympus, unfocused, 28.5 mm face diameter) were mounted to a breadboard such that they faced each other. One transducer served as a transmitter and one as a receiver. The distance between the transducers’ faces was 100 mm. The breadboard with the mounted transducers was positioned at the bottom of the water tank. The center of each face of the transducer was 145 mm above the breadboard. Each skull was electronically translated and rotated such that 1) the line connecting the centers of the two transducers intersected with the 4 markers 2) all segments of the skull were in the far field of the transmitting transducer (at a distance greater than 68.6 mm). The line connecting the centers of the two transducers was implemented using a custom 3D printed plastic pointer that was positioned into the holder of the transmitting transducer.

### 2.5. Ultrasound system and pulses

The through-transmit protocol was implemented on the Vantage256 system (Verasonics), using a custom matlab script. We used chirp pulses of 3 distinct forms. Chirps are frequency-modulated waveforms that have a narrow autocorrelation function. A narrow autocorrelation function maximizes the accuracy of the detection of time delays in through-transmit procedures [54, 55]. Chirp3 consistent of three consecutive cycles of [0.75, 1, 1.25] times the center frequency of 500 kHz. Chirp4 consistent of four consecutive cycles with [0.75, 1, 1.25, 1.5] times the center frequency. Chirp5 consistent of five consecutive cycles with [0.5, 0.75, 1, 1.25, 1.5] times the center frequency. Our transducers were broadband (Videoscan series, Olympus) and so capable of emitting the frequency spectrum. Each pulse was transmitted and detected 32 times; the responses were averaged together.

### 2.6. Through-transmit procedure

The through-transmit procedure quantifies the changes in amplitudes and received times following an introduction of an object (e.g., skull) into the transmit-receive path. Data were first collected in water. This provided an average no-skull receive waveform. Consequently, the skull was lowered into a location described above, and gradually rotated in 1 degree increments, with the through-transmit data taken at each. This provided through-skull receive waveforms. The maximum of each through-skull receive waveform divided by the maximum of the no-skull receive waveform provided the relative pressure transmission, separately for each angle (Fig. 3, Fig. 5). To determine the time shift, a cross-correlation was computed between the no-skull and through-skull receive waveforms, separately for each angle. The peak of the cross-correlation defined the time shift. This time corresponds to the speedup values shown in Fig. 6 and Fig. 7. The three pulses produced equivalent values of time shift in most cases. These values were averaged together.

**Figure 3:**
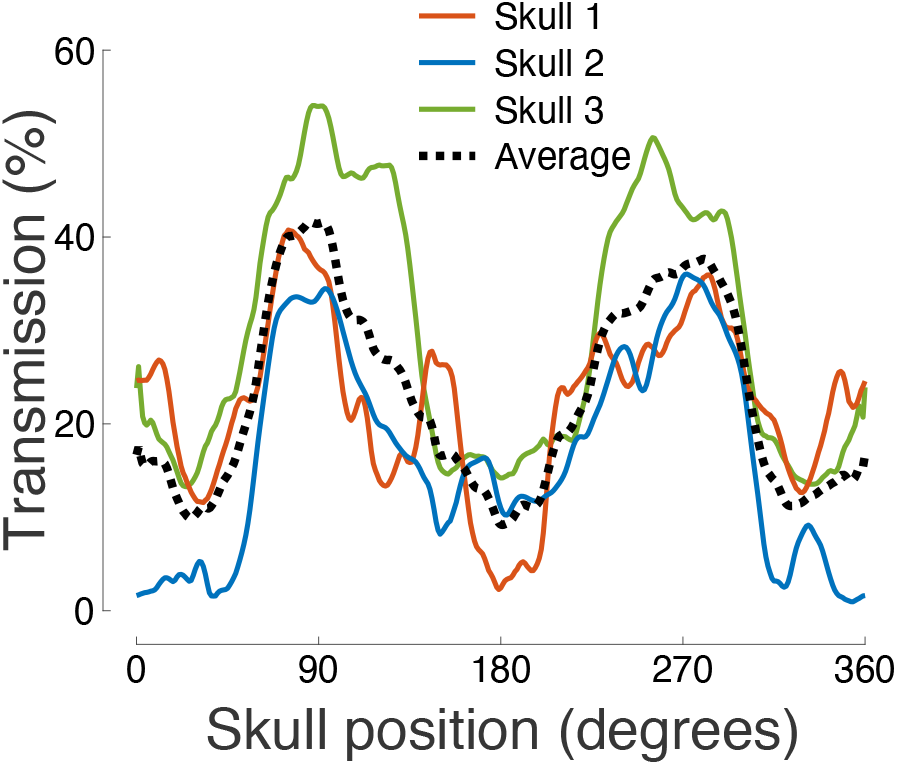
Ultrasound transmission throughout the skull. The figure shows the relative pressure attenuation (skull versus no skull) for each measured segment of the skull. The carrier frequency was 500 kHz.

**Figure 4:**
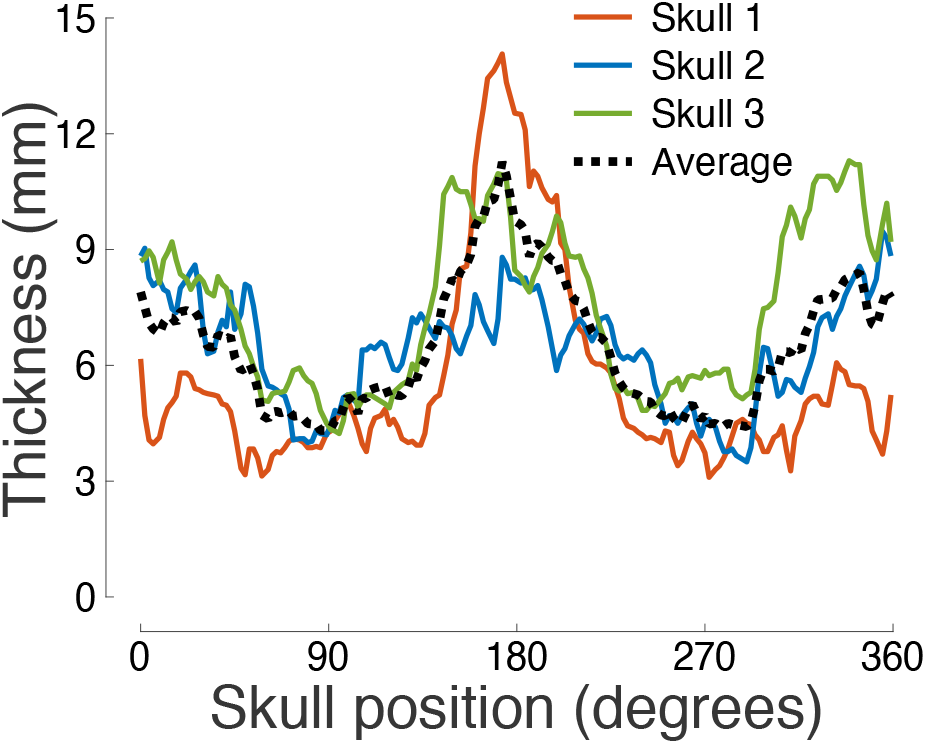
Skull thickness across angular position. Mean thickness of each segment of the skull within the through-transmit plane. Three independent measurements were taken at each position.

**Figure 5:**
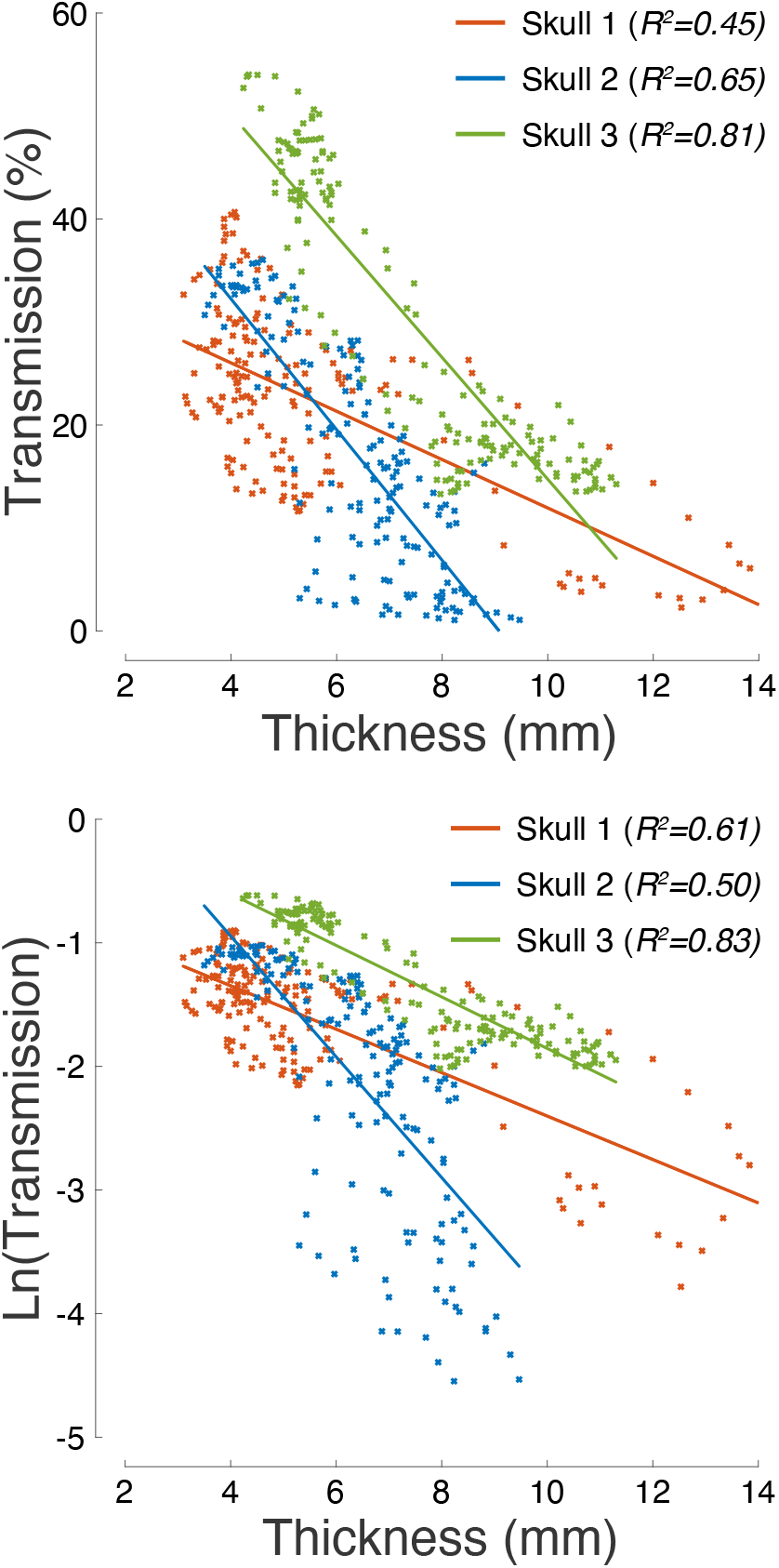
Ultrasound transmission is strongly governed by skull thickness. Ultrasound pressure transmission (top) and its natural logarithm (bottom) as a function of skull thickness. The *R*^2^ values listed in the inset provide the amount of variance explained by the linear fits superimposed on the plots.

**Figure 6:**
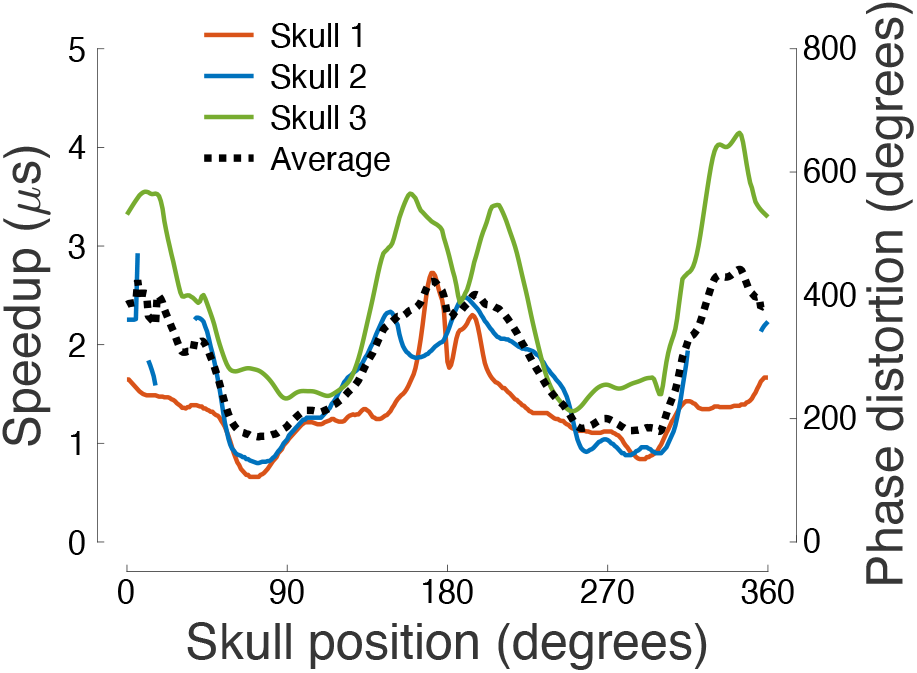
Speedup and phase distortion across the skull. Ultrasound speedup through the skull (*τ*) and the associated phase distortion (*ωτ*) as a function of the skull position. Several segments of the occipital and frontal bones in Skull 2 provided extreme aberration, rendering the through-transmit cross-correlation unreliable; values for these segments are therefore not shown.

**Figure 7:**
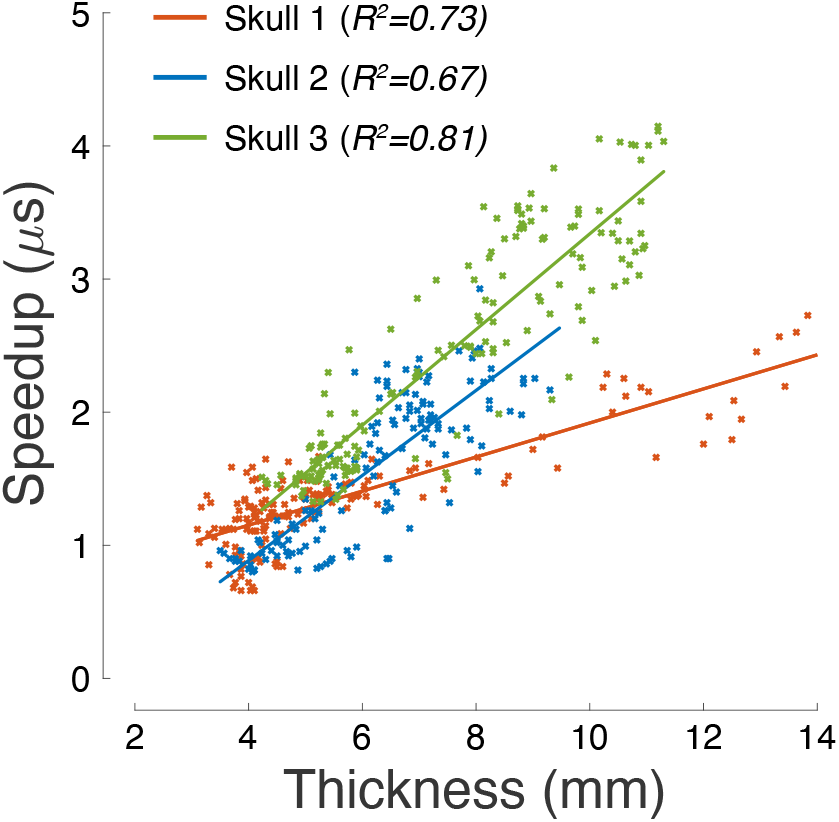
Ultrasound phase distortion is proportional to skull thickness. Ultrasound speedup through individual segments of the skull as a function of skull thickness. The *R*^2^ values listed in the inset provide the amount of variance explained by the linear fits superimposed on the plots.

### 2.7. Through-transmit relationships

The ultrasound speedup through a skull segment *τ* is mathematically proportional to the thickness of the segment *h* through:

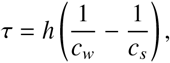

where *c_w_* = 1481 m/s is the speed of sound through water, and *c_s_* is the average speed of sound through the skull segment. The phase distortion or shift caused by the speedup is equal to *ωτ*, where *ω* is the angular frequency.

The measured pressure transmission (*T*) depends on ultrasound reflection from the individual layers of the skull and on a set of attenuation factors (*α*) that include ultrasound absorption and scattering. Under an assumption that these factors can be considered separate and independent, we can write:

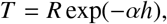

where *R* is the loss due to reflection and *h* is the skull segment thickness. Taking a logarithm of this equation yields:

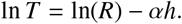

The set of thickness-dependent attenuation factors *α* can be inferred as a slope of the relationship between ln(*T*) and *h* (Fig. 5).

## 3. Results

We assessed the acoustic properties across intact *ex-vivo* human skulls using an apparatus that allowed controlled rotation about a central axis (Fig. 1). The acoustic properties were characterized using standard through-transmit measurements (Materials and Methods), with the transmitting/receiving transducer positioned outside/inside of the skull. The position of the two transducers was fixed; only the skull was rotated. The through-transmit values obtained through the skull were compared to free-field values in which no skull was present, as in previous studies [37, 51].

We performed the through-transmit measurements in three degassed and hydrated *ex-vivo* skulls. The CT images of the skulls within the through-transmit plane are shown in Fig. 2.

The relative pressure transmission across the individual anatomical locations is shown in (Fig. 3). The figure reveals that the highest transmission was observed in segments centered over the parietal bone. The average transmission in the parietal regions was 2.4 higher than in the temporal regions and 2.0 higher than in the occipital regions (Table 1). The transmission was maximal within the parietal bone and delivered an average 41.6% of the ultrasound pressure across the three skulls. As apparent from the figure, the transmission varied substantially across the 3 subjects—the 3 skulls showed an average 22.2%, 16.8%, and 29.5% transmission, respectively. Effects of the skull position and subject were both significant. In particular, a two-way ANOVA revealed a significant modulation of the pressure transmission by the skull position (*F*(360, 720) = 7.48, *p* = 4.7 × 10^−115^) and by subject (*F*(2, 720) = 353.90, *p* = 9.1 × 10^−108^).

**Table 1:**
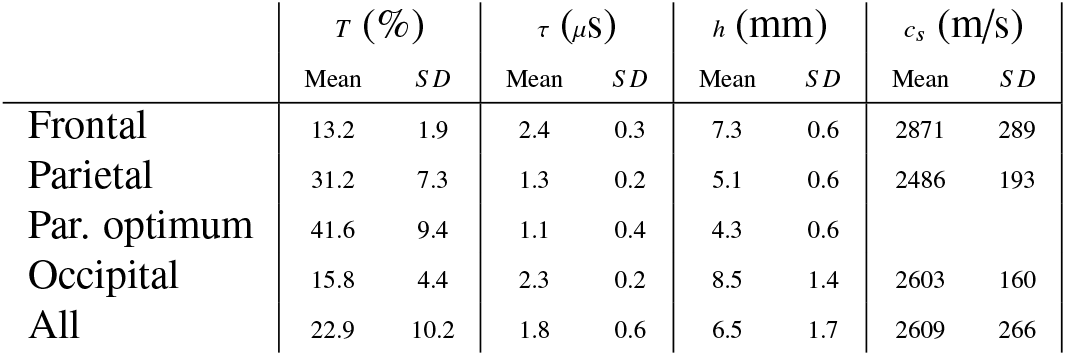
Acoustic properties quantified over distinct skull regions. The table lists the mean±SD transmission (*T*), speedup (*τ*), thickness (*h*), and speed of sound (*c_s_*) as a function of skull position (rows). The parietal optimum entry lists maximal transmission, minimal speedup, and minimal thickness, averaged across the 3 skulls.

We measured the thickness of the skulls across the through-transmit plane using a caliper (see Materials and Methods). The skull thickness as a function of position is shown in Fig. 4. The figure reveals a substantial variability in skull thickness within and across individuals. The thickness ranged from 3.1 mm to 14.0 mm, with an average of 6.5 mm and a standard deviation of 1.7 mm. The parietal bone was on average thinner than the frontal and occipital bones (Table 1). The thickness varied across the 3 subjects, with average values of 5.6 mm, 6.4 mm, and 7.5 mm, respectively. A two-way ANOVA revealed a significant modulation of thickness by the skull position (*F*(66, 332) = 3.75, *p* = 7.2 × 10^−25^) and by subject (*F*(2, 332) = 64.96, *p* = 1.6 × 10^−24^).

The profiles of pressure transmission (Fig. 3) and skull thickness (Fig. 4) suggest an inverse relationship between these quantities: the thinner the skull, the more effective the transmission. Indeed, we found that skull thickness is a strong predictor of the transmission efficacy (Fig. 5). In the three skulls, skull thickness explained 45%, 65%, and 81% of the variance in the pressure transmission, respectively. An ANCOVA model, with a continuous factor of skull thickness and a discrete factor of subject, found a significant effect of thickness (*F*(1, 495) = 974.45, *p* = 4.9 × 10^−119^) as well as an effect of subject (*F*(2, 495) = 161.15, *p* = 1.3 × 10^−54^) and thickness×subject interaction (*F*(2, 495) = 90.31, *p* = 3.7 × 10^−34^). Thus, there is a significant difference in the pressure transmission across the subjects, and the dependence of transmission on skull thickness is subject-specific. We further investigated the thickness dependence of the natural logarithm of the pressure transmission. The logarithmic formulation may enable the quantification of skull thickness-dependent attenuation factors *α* from the slope of the linear relationships showed in the bottom part of Fig. 5 (see Materials and Methods). The slopes for the 3 skulls were −0.176, −0.489, and −0.208, respectively. According to the simple—though likely simplistic—model (Materials and Methods), this slope translates into thickness-dependent attenuation factors of *α* = 176 Np/m, *α* = 489 Np/m, and *α* = 208 Np/m, respectively.

It is known that the skull speeds up the propagation of ultrasound compared to water. This relative speedup leads to distortions or shifts of the ultrasound phase. We investigated how this speedup and phase distortion depend on the individual segments of the skull. We found that the distortion was smallest for the parietal regions (Fig. 6), with a value 1.8 times smaller than for the frontal and 1.7 times smaller than the occipital regions (Table 1). A two-way ANOVA revealed a significant modulation of the phase distortion by the skull position (*F*(360, 654) = 5.08, *p* = 1.5 × 10^−72^) and by subject (*F*(2, 654) = 694.63, *p* = 1.7 × 10^−162^).

The speedup and the associated phase distortion should be linearly proportional to skull thickness (see Materials and Methods). This proportionality has been demonstrated previously [45]. Indeed, we confirmed these findings (Fig. 7). In the 3 skulls, skull thickness explained 73%, 78%, and 81% of the variance in the phase aberration, respectively. An ANCOVA model, again with a continuous factor of thickness and a discrete factor of subject, found a significant effect of thickness (*F*(1, 464) = 1175.53, *p* = 2.9 × 10^−129^) as well as an effect of subject (*F*(2, 464) = 55.92, *p* = 1.7 × 10^−22^) and thickness×subject interaction (*F*(2, 464) = 142.78, *p =* 4.8 × 10^−49^). Thus, the linear relationship between the phase aberration and skull thickness, albeit robust within an individual, is variable across individuals.

The previous finding suggests an appreciable difference in the average speed of sound across the skulls. We investigated the speed of sound across the skulls and across the individual segments of each skull (Fig. 8). There was a substantial variability in the speed of sound, both across and within the subjects. In particular, the three skulls showed a mean SD of 2451 383 m/s, 2401±307 m/s, and 2887±412 m/s, respectively. Although there was a trend for the speed of sound to be lower for the parietal bone (Table 1) compared to frontal and occipital bone, this was not consistent across subjects (Fig. 8). A two-way ANOVA confirmed the significant variability of the speed of sound by skull position (*F*(359, 652) = 1.29, *p* = 0.0024) and by subject (*F*(2, 652) = 173.07, *p* = 5.1 × 10^−61^).

**Figure 8:**
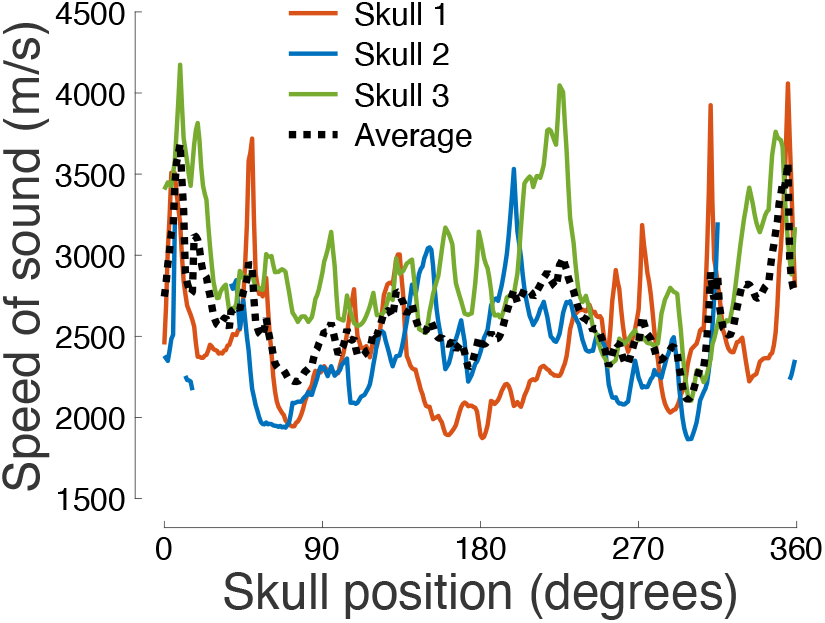
Speed of sound across the skull. The speed of sound (*c_s_*) determined from the through-transmit *τ* values (see Materials and Methods) as a function of the skull position.

## 4. Discussion

In this study, we investigated the transmission and phase distortion of ultrasound across intact human skulls.

We found that 500 kHz ultrasound—a common frequency for transcranial applications in humans—was most effectively transmitted through segments of the parietal bone (Fig. 3), in all three *ex-vivo* specimens. We further found that the transmission was strongly dependent on the skull thickness: the thinner the skull, the more effective the transmission (Fig. 5). Since parietal bone was found to be the thinnest on average (Fig. 4, Table 1), this negative correlation can partially explain the facilitated transmission through the parietal bone.

A negative correlation between skull thickness and ultrasound transmission could be expected, but has not, to our knowledge, been shown explicitly. Our direct measurements of skull thickness using a caliper uncovered a surprisingly tight relationship (64% of variance explained, mean across 3 skulls) between the skull thickness and ultrasound transmission. However, the slopes of the dependence on the thickness varied across the 3 skulls. In addition—and as observed in previous studies [37, 39, 40]—we found a significant difference in the average transmission across the 3 skulls. These subject-specific factors complicate a parsimonious description of ultrasound transmission based on skull thickness alone.

We found that the speedup of ultrasound due to the propagation through the skull bone, and the associated phase distortion, are also smallest over parietal bone segments (Fig. 6). This finding can be explained by our (Fig. 7) and previous [45] findings that the speedup and phase shift are proportional to skull thickness. The slope of this dependence, which is a function of the average speed of sound through the skull (see Materials and Methods), varied significantly across the skulls. This variability has been observed also in previous studies, but can be accounted for using CT skull density measurements [51, 56].

The data provided in this study corroborate the notion of substantial variability of acoustic properties across individuals [29, 37–40]. In addition to the inter-subject variability, we have shown, in a systematic experiment, that there is substantial skull location-dependent variability. Transmission and phase shift comprised a well-defined function of skull location (Fig. 3, Fig. 6). The speed of sound, in contrast, showed a much less predictable pattern (Fig. 8).

The primary goal of our study was to identify the regions of the skull that provide optimal transmission. Although our robotic system allowed us to assess the acoustic properties across the perimeter of the skull, it only enabled us to operate along the middle segments of the skull. Acoustic properties for more dorsal portion of calvariae are to be found elsewhere [38, 39]. It should also be noted that *ex-vivo* skulls do not model fresh or well-preserved skulls exactly [37]. Therefore, the values presented in this study should be taken as relative—with respect to anatomical location or skull thickness, rather than absolute.

In this study, we could directly measure skull thickness using a caliper to within 0.1 mm precision. Using these measurements, we uncovered a surprisingly tight relationship between ultrasound transmission and skull thickness. This relationship invites future investigations with the goal to account for the attenuation of ultrasound transmission through the skull based on CT [57], MRI [57, 58], or other imaging modalities [53].

In summary, we measured the acoustic properties throughout the perimeter of intact human *ex-vivo* skulls. We found that regions of the parietal bone provide much more effective transmission and lower phase distortion than frontal and occipital regions. We further found that the ultrasound transmission strongly depends on the skull thickness. These data inform future approaches to compensate for ultrasound skull aberrations and guide the design of future devices for safe, effective, and reproducible transcranial ultrasound therapies.

## 5. Acknowledgements

This work was supported by the National Institutes of Health (grants R00NS100986 and F32MH123019).

## References

[1] K. Hynynen, F. A. Jolesz, Demonstration of potential noninvasive ultrasound brain therapy through an intact skull, Ultrasound in medicine & biology 24 (2) (1998) 275–283.

[2] E. Landhuis, Ultrasound for the brain, Nature 551 (7679) (2017) 257–259.

[3] Y. Meng, K. Hynynen, N. Lipsman, Applications of focused ultrasound in the brain: From thermoablation to drug delivery, Nature Reviews Neurology (2020) 1–16.

[4] P. Ghanouni, K. B. Pauly, W. J. Elias, J. Henderson, J. Sheehan, S. Monteith, M. Wintermark, Transcranial MRI-guided focused ultrasound: a review of the technologic and neurologic applications, American Journal of Roentgenology 205 (1) (2015) 150–159.

[5] M. Giordano, V. M. Caccavella, I. Zaed, L. F. Manzillo, N. Montano, A. Olivi, F. M. Polli, Comparison between deep brain stimulation and magnetic resonance-guided focused ultrasound in the treatment of essential tremor: a systematic review and pooled analysis of functional outcomes, Journal of Neurology, Neurosurgery & Psychiatry 91 (12) (2020) 1270–1278.

[6] A. Carpentier, M. Canney, A. Vignot, V. Reina, K. Beccaria, C. Horodyckid, C. Karachi, D. Leclercq, C. Lafon, J.-Y. Chapelon, et al., Clinical trial of blood-brain barrier disruption by pulsed ultrasound, Science Transl. Medicine 8 (343) (2016) 343.

[7] N. Lipsman, Y. Meng, A. J. Bethune, Y. Huang, B. Lam, M. Masellis, N. Herrmann, C. Heyn, I. Aubert, A. Boutet, et al., Blood–brain barrier opening in Alzheimer’s disease using MR-guided focused ultrasound, Nature Communications 9 (1) (2018) 2336.

[8] P. Anastasiadis, J. A. Winkles, A. J. Kim, G. F. Woodworth, Focused ultrasound-mediated blood-brain barrier disruption for enhanced drug delivery to brain tumors, in: Nanotherapy for Brain Tumor Drug Delivery, Springer, 2021, pp. 205–223.

[9] R. D. Airan, R. A. Meyer, N. P. Ellens, K. R. Rhodes, K. Farahani, M. G. Pomper, S. D. Kadam, J. J. Green, Noninvasive targeted transcranial neuro-modulation via focused ultrasound gated drug release from nanoemulsions, Nano Letters 17 (2) (2017) 652–659.

[10] J. B. Wang, M. Aryal, Q. Zhong, D. B. Vyas, R. D. Airan, Noninvasive ultrasonic drug uncaging maps whole-brain functional networks, Neuron 100 (3) (2018) 728–738.

[11] H. Lea-Banks, Y. Meng, S.-K. Wu, R. Belhadjhamida, C. Hamani, K. Hynynen, Ultrasound-sensitive nanodroplets achieve targeted neuromodulation, Journal of Controlled Release (2021).

[12] J. Kubanek, Neuromodulation with transcranial focused ultrasound, Neurosurgical Focus 44 (2) (2018) E14.

[13] W. J. Tyler, S. W. Lani, G. M. Hwang, Ultrasonic modulation of neural circuit activity, Current opinion in neurobiology 50 (2018) 222–231.

[14] J. Blackmore, S. Shrivastava, J. Sallet, C. R. Butler, R. O. Cleveland, Ultrasound neuromodulation: A review of results, mechanisms and safety, Ultrasound in medicine & biology 45 (7) (2019) 1509–1536.

[15] J. Kubanek, J. Brown, P. Ye, K. B. Pauly, T. Moore, W. Newsome, Remote, brain region–specific control of choice behavior with ultrasonic waves, Science Advances 6 (21) (2020) eaaz4193.

[16] J. L. Sanguinetti, S. Hameroff, E. E. Smith, T. Sato, C. M. Daft, W. J. Tyler, J. J. Allen, Transcranial focused ultrasound to the right prefrontal cortex improves mood and alters functional connectivity in humans, Frontiers in Human Neuroscience 14 (2020) 52.

[17] B. W. Badran, K. A. Caulfield, S. Stomberg-Firestein, P. M. Summers, T. Dowdle, M. Savoca, X. Li, C. W. Austelle, E. B. Short, J. J. Borckardt, et al., Sonication of the anterior thalamus with mri-guided transcranial focused ultrasound (tfus) alters pain thresholds in healthy adults: A double-blind, sham-controlled study, Brain Stimulation 13 (6) (2020) 1805–1812.

[18] L. Verhagen, C. Gallea, D. Folloni, C. Constans, D. E. Jensen, H. Ahnine, L. Roumazeilles, M. Santin, B. Ahmed, S. Lehéricy, et al., Offline impact of transcranial focused ultrasound on cortical activation in primates, Elife 8 (2019) e40541.

[19] E. F. Fouragnan, B. K. Chau, D. Folloni, N. Kolling, L. Verhagen, M. Klein-Flügge, L. Tankelevitch, G. K. Papageorgiou, J.-F. Aubry, J. Sallet, et al., The macaque anterior cingulate cortex translates counterfactual choice value into actual behavioral change, Nature neuroscience 22 (5) (2019) 797–808.

[20] N. Khalighinejad, A. Bongioanni, L. Verhagen, D. Folloni, D. Attali, J.-F. Aubry, J. Sallet, M. F. Rushworth, A basal forebrain-cingulate circuit in macaques decides it is time to act, Neuron 105 (2) (2020) 370–384.

[21] V. Velling, S. Shklyaruk, Modulation of the functional state of the brain with the aid of focused ultrasonic action, Neuroscience and behavioral physiology 18 (5) (1988) 369–375.

[22] R. F. Dallapiazza, K. F. Timbie, S. Holmberg, J. Gatesman, M. B. Lopes, J. Price, G. W. Miller, W. J. Elias, Noninvasive neuromodulation and thalamic mapping with low-intensity focused ultrasound, Journal of Neurosurgery (2017) 1–10.

[23] W. S. Chang, H. H. Jung, E. Zadicario, I. Rachmilevitch, T. Tlusty, S. Vitek, J. W. Chang, Factors associated with successful magnetic resonance-guided focused ultrasound treatment: efficiency of acoustic energy delivery through the skull, Journal of neurosurgery 124 (2) (2016) 411–416.

[24] J. K. Mueller, L. Ai, P. Bansal, W. Legon, Numerical evaluation of the skull for human neuromodulation with transcranial focused ultrasound, Journal of neural engineering 14 (6) (2017) 066012.

[25] H. Chen, E. E. Konofagou, The size of blood–brain barrier opening induced by focused ultrasound is dictated by the acoustic pressure, Journal of Cerebral Blood Flow & Metabolism 34 (7) (2014) 1197–1204.

[26] W. Lee, H.-C. Kim, Y. Jung, Y. A. Chung, I.-U. Song, J.-H. Lee, S.- S. Yoo, Transcranial focused ultrasound stimulation of human primary visual cortex, Scientific Reports 6 (2016).

[27] V. Braun, J. Blackmore, R. O. Cleveland, C. R. Butler, Transcranial ultrasound stimulation in humans is associated with an auditory confound that can be effectively masked, Brain stimulation 13 (6) (2020) 1527–1534.

[28] M. Plaksin, E. Kimmel, S. Shoham, Cell-type-selective effects of intramembrane cavitation as a unifying theoretical framework for ultrasonic neuromodulation, eneuro 3 (3) (2016).

[29] A. Y. Ammi, T. D. Mast, I.-H. Huang, T. A. Abruzzo, C.-C. Coussios, G. J. Shaw, C. K. Holland, Characterization of ultrasound propagation through ex-vivo human temporal bone, Ultrasound in medicine & biology 34 (10) (2008) 1578–1589.

[30] S. Purkayastha, F. Sorond, Transcranial doppler ultrasound: technique and application, in: Seminars in neurology, Vol. 32, NIH Public Access, 2012, p. 411.

[31] Z. Wang, T. Komatsu, H. Mitsumura, N. Nakata, T. Ogawa, Y. Iguchi, M. Yokoyama, An uncovered risk factor of sonothrombolysis: substantial fluctuation of ultrasound transmittance through the human skull, Ultrasonics 77 (2017) 168–175.

[32] D. E. Soulioti, D. Espíndola, P. A. Dayton, G. F. Pinton, Super-resolution imaging through the human skull, IEEE transactions on ultrasonics, ferroelectrics, and frequency control 67 (1) (2019) 25–36.

[33] W. Legon, T. F. Sato, A. Opitz, J. Mueller, A. Barbour, A. Williams, W. J. Tyler, Transcranial focused ultrasound modulates the activity of primary somatosensory cortex in humans, Nature Neuroscience 17 (2) (2014) 322–329.

[34] W. Lee, H. Kim, Y. Jung, I.-U. Song, Y. A. Chung, S.-S. Yoo, Imageguided transcranial focused ultrasound stimulates human primary somatosensory cortex, Scientific reports 5 (2015).

[35] W. Legon, L. Ai, P. Bansal, J. K. Mueller, Neuromodulation with single element transcranial focused ultrasound in human thalamus, Human brain mapping 39 (5) (2018) 1995–2006.

[36] W. Legon, P. Bansal, R. Tyshynsky, L. Ai, J. K. Mueller, Transcranial focused ultrasound neuromodulation of the human primary motor cortex, Scientific reports 8 (1) (2018) 1–14.

[37] F. J. Fry, J. E. Barger, Acoustical properties of the human skull, The Journal of the Acoustical Society of America 63 (5) (1978) 1576–1590.

[38] P. J. White, G. T. Clement, K. Hynynen, Longitudinal and shear mode ultrasound propagation in human skull bone, Ultrasound in medicine & biology 32 (7) (2006) 1085–1096.

[39] S. Pichardo, V. W. Sin, K. Hynynen, Multi-frequency characterization of the speed of sound and attenuation coefficient for longitudinal transmission of freshly excised human skulls, Physics in Medicine & Biology 56 (1) (2010) 219.

[40] S. Pichardo, C. Moreno-Hernández, R. A. Drainville, V. Sin, L. Curiel, K. Hynynen, A viscoelastic model for the prediction of transcranial ultrasound propagation: application for the estimation of shear acoustic properties in the human skull, Physics in Medicine & Biology 62 (17) (2017) 6938.

[41] F. Fry, Transkull transmission of an intense focused ultrasonic beam, Ultrasound in medicine & biology 3 (2-3) (1977) 179–184.

[42] J. E. Barger, Attenuation and dispersion of ultrasound in cancellous bone, Ultrasonic Tissue Characterization 11 (1979) 197–201.

[43] J. Evans, M. Tavakoli, Ultrasonic attenuation and velocity in bone, Physics in Medicine & Biology 35 (10) (1990) 1387.

[44] M. Tavakoli, J. Evans, Dependence of the velocity and attenuation of ultrasound in bone on the mineral content, Physics in Medicine & Biology 36 (11) (1991) 1529.

[45] G. Clement, K. Hynynen, Correlation of ultrasound phase with physical skull properties, Ultrasound in medicine & biology 28 (5) (2002) 617–624.

[46] G. Pinton, J.-F. Aubry, E. Bossy, M. Muller, M. Pernot, M. Tanter, Attenuation, scattering, and absorption of ultrasound in the skull bone, Medical physics 39 (1) (2012) 299–307.

[47] K. Hynynen, J. Sun, Trans-skull ultrasound therapy: the feasibility of using image-derived skull thickness information to correct the phase distortion, IEEE transactions on ultrasonics, ferroelectrics, and frequency control 46 (3) (1999) 752–755.

[48] J. Aarnio, G. T. Clement, K. Hynynen, A new ultrasound method for determining the acoustic phase shifts caused by the skull bone, Ultrasound in medicine & biology 31 (6) (2005) 771–780.

[49] P. White, G. Clement, K. Hynynen, Local frequency dependence in transcranial ultrasound transmission, Physics in Medicine & Biology 51 (9) (2006) 2293.

[50] L. Marsac, D. Chauvet, R. La Greca, A.-L. Boch, K. Chaumoitre, M. Tanter, J.-F. Aubry, Ex vivo optimisation of a heterogeneous speed of sound model of the human skull for non-invasive transcranial focused ultrasound at 1 mhz, International Journal of Hyperthermia 33 (6) (2017) 635–645.

[51] T. D. Webb, S. A. Leung, J. Rosenberg, P. Ghanouni, J. J. Dahl, N. J. Pelc, K. B. Pauly, Measurements of the relationship between ct hounsfield units and acoustic velocity and how it changes with photon energy and reconstruction method, IEEE transactions on ultrasonics, ferroelectrics, and frequency control 65 (7) (2018) 1111–1124.

[52] S. A. Leung, T. D. Webb, R. R. Bitton, P. Ghanouni, K. B. Pauly, A rapid beam simulation framework for transcranial focused ultrasound, Scientific reports 9 (1) (2019) 1–11.

[53] L. Deng, A. Hughes, K. Hynynen, A noninvasive ultrasound resonance method for detecting skull induced phase shifts may provide a signal for adaptive focusing, IEEE Transactions on Biomedical Engineering 67 (9) (2020) 2628–2637.

[54] M. Crocco, P. Pellegretti, C. Sciallero, A. Trucco, Combining multipulse excitation and chirp coding in contrast-enhanced ultrasound imaging, Measurement Science and Technology 20 (10) (2009) 104017.

[55] S. Callegari, M. Ricci, S. Caporale, M. Monticelli, M. Eroli, L. Senni, R. Rovatti, G. Setti, P. Burrascano, From chirps to random-fm excitations in pulse compression ultrasound systems, in: 2012 IEEE International Ultrasonics Symposium, IEEE, 2012, pp. 471–474.

[56] J.-F. Aubry, M. Tanter, M. Pernot, J.-L. Thomas, M. Fink, Experimental demonstration of noninvasive transskull adaptive focusing based on prior computed tomography scans, The Journal of the Acoustical Society of America 113 (1) (2003) 84–93.

[57] T. D. Webb, S. A. Leung, P. Ghanouni, J. J. Dahl, N. J. Pelc, K. B. Pauly, Acoustic attenuation: Multifrequency measurement and relationship to ct and mr imaging, IEEE Transactions on Ultrasonics, Ferroelectrics, and Frequency Control (2020) 1–1 doi:10.1109/TUFFC.2020.3039743.

[58] G. W. Miller, M. Eames, J. Snell, J.-F. Aubry, Ultrashort echo-time mri versus ct for skull aberration correction in mr-guided transcranial focused ultrasound: In vitro comparison on human calvaria, Medical physics 42 (5) (2015) 2223–2233.

